# Phytorisque: an Integrated Assessment Tool for Evaluating the Environmental Risk of Pesticides

**DOI:** 10.64898/2026.04.30.721842

**Authors:** Loïc Monseur, Jean-Baptiste de Maere, Chloé Guillitte, Gaspard Nihorimbere, Laurence Janssens, Claude Bragard

**Affiliations:** Coordination recherche et développement rural (CORDER), Louvain-la-Neuve, Belgium; Université catholique de Louvain, Earth & Life Institute, Louvain-la-Neuve, Belgium

**Keywords:** Ecotoxicity, Bioaccumulation, Mobility, Persistence, Leaching, Runoff, Biodegradation, Modelling, Volatilization, Active Substance

## Abstract

**Introduction:** The environmental impacts of pesticides have raised increasing concern, prompting the development of indicators to assess associated risks across ecosystems. Two main categories are generally distinguished: score-based indicators, which aggregate variables into scores, and risk-based indicators, grounded in the definition of risk as the product of hazard and exposure. Although more data-intensive and more complex to implement, risk-based indicators are recognized to better preserve proportionality with actual risk levels.

**Objectives:** This study presents Phytorisque, a model based on the exposure–toxicity ratio to monitor risks associated with pesticide use in Walloon agriculture, from farm to regional scales, and to identify the most contributing active substances in support of risk-reduction policies

**Method:** Phytorisque is a hybrid model that combines mechanistic, empirical, and statistical approaches, integrating quantities of active substances, their ecotoxicological characteristics, and their mobility, persistence, and bioaccumulation properties to generate indices specific to different environmental compartments.

**Results:** The indices obtained enable comparison across substances, agricultural sectors, years, and management scenarios. The Phytorisque model provides an integrated assessment of risk across environmental compartments. It can monitor risk evolution over the years for policy impacts evaluation, diagnose the most problematic substances and prospect environmental risks associated with the use of chemical phytoproducts.

**Conclusions:** Phytorisque provides an integrated risk assessment approach adapted to temporal monitoring, diagnosis, and forecasting. It is a relevant operational tool for supporting regional strategies aimed at reducing pesticide-related risks. The model is also transferable to other regions through the adaptation of parameters to local conditions and context.

## 1. INTRODUCTION

The use of phytopharmaceutical products (PPPs) in agriculture has raised increasing concerns regarding their impact on environmental ecosystems. Numerous negative effects of pesticide use were reported, including damage to groundwater or terrestrial and aquatic ecosystems (Vercruysse et al., 2002; Gensch et al., 2024; Köninger et al., 2026; Wolfram et al., 2026). Reducing these impacts has become a major objective of agricultural and environmental policies worldwide.

In the European Union, the Sustainable Use of Pesticides Directive (2009/128/EC) was adopted, requiring Member States to establish National Action Plans to reduce both the quantity of pesticides used and the risks associated with their use. Several indicators were subsequently developed to evaluate reductions in pesticide use and associated risks.

According to the Organisation for Economic Cooperation and Development (OCDE, 1998), the pesticide indicators are classified in two main categories: scoring indicators (SI) and exposure-toxicity ratio indicators (ETR). SI convert input variables, such as physicochemical properties, ecotoxicological data, or formulation characteristics, into an aggregated final score. Examples include the Pesticide Load Indicator (Kudsk et al., 2018) and the Environmental Impact Quotient (Obregon et al., 2025). In contrast, ETR-based indicators rely on the definition of risk, expressed as the ratio between exposure and toxicity. Indicators such as SYNOPS and Environmental Yardstick for Pesticides are consistent with this definition (Reus et al., 2000; Strassemeyer et al., 2017). SI are straightforward to implement across different contexts, but the aggregation of scores complexifies results interpretations, and scores variation are not necessarily proportional to risk evolution for the environment (OCDE, 1998; Reus et al., 2002). Conversely, ETR indicators generally provide a more robust interpretation of the risk. Their application, however, is often constrained by their inherent complexity and data requirements needed to accurately represent pesticide fate and behavior in their specific region of application (Reus et al., 2002; Kudsk et al., 2018; Pierlot et al., 2023).

Most pesticide risk indicators are location-specific and were developed to address specific objectives at the regional or national level. In the Walloon Region, the Pesticide Reduction Program (Programme Wallon de Réduction des Pesticides, PWRP) provides the framework for reducing the environmental impact of pesticide use (PWRP III 2023-2027). At present, the monitoring of pesticide use is mainly based on quantitative indicators (Corder ASBL, 2025a), without considering the environmental fate or the toxicity to non-target organisms in ecosystems. Therefore, there is a need for a recognized tool able to assess risk trends across environmental compartments such as soil, surface water, and groundwater at the regional scale.

To address this objective, the Phytorisque was developed as an environmental risk assessment model. It generates integrated indices that reflect the risk associated with the use of PPP’s active substances (a.s.) in agriculture for different environmental compartments within the Walloon territory. The model incorporates the diversity of processes governing environmental fate (transfer, dissipation, and accumulation), as well as toxicity to non-target organisms. It also accounts for the hydropedological and meteorological conditions specific to Wallonia.

Phytorisque is intended to evaluate the impact of risk reduction strategies by monitoring the evolution of risk associated with a.s. uses in agriculture. In addition, the tool is designed to assess the risks associated with a.s. used in arable crops across environmental compartments and can be used for prospective scenario analysis. The model is conceived as a flexible and evolving framework, with the potential to be adapted to other regions through the adjustment of parameters to local pedoclimatic and land-use conditions.

## 2. MATERIALS AND METHOD

### 2.1. Data sources

#### 2.1.1. Meteorological, hydrological and pedological data

Meteorological and hydropedological data specific to Wallonia were collected from several regional sources: the Wallonia Geoportal (Département Données transversales (SPW), October-1-2025) for soil and hydrological data as well as agricultural land parcels, and the Agromet network of the Walloon Agricultural Research Centre (Rosillon et al., 2024) for meteorological data.

To calibrate the model for conditions representative of arable crops in Wallonia, average soil and hydrological parameters were calculated using the QGIS software. After retrieving spatial raster layers describing soil and hydrological properties at a 5 m resolution, these layers were clipped using masks identifying areas of conventional arable crops. Subsequently, the mean or median pixel value, depending on the data collected, were calculated at the Walloon scale to obtain representative environmental parameter values. Meteorological parameters were extracted from data recorded at the Sombreffe station of the Agromet network, which is centrally located with respect to the main arable cropping areas in Wallonia.

#### 2.1.2. Active substances properties and quantities

To illustrate outputs generated by the model, a focus was made on five a.s. reflecting the physicochemical properties diversity of substances used in arable cropping systems in Wallonia. Physicochemical and ecotoxicological properties were obtained from the Pesticides Properties DataBase (PPDB) (Lewis et al., 2016). Quantity of a.s. used come from a regional study that estimates a.s. uses in 15 agricultural sectors representative of Walloon arable farming systems, accounting for 92% of the conventional arable crop area in Wallonia. They are established by a statistical extrapolation method from a sample of farms, using accounting data provided by the Direction de l’Analyse Economique Agricole (DAEA). The methodology and results are detailed in Corder (2025a). Physicochemical properties and quantities used in 2022 of these 5 a.s. are presented in Table 1.

**Table 1:**
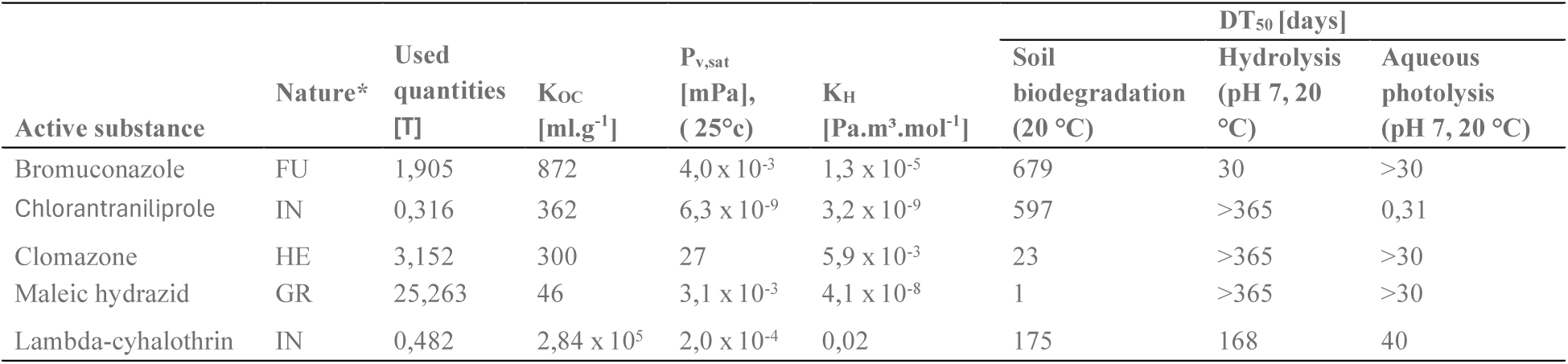
Estimated quantities of active substances used in conventional arable farming in Wallonia in 2022 (Corder ASBL, 2025a) and physicochemical properties of these substances (Lewis et al., 2016). * FU: Fungicide; IN: Insecticide; HE: Herbicide; GR: Growth regulator, T: toxicity factor, KOC: organic carbon partition coefficient, Pv,sat: vapour pressure, KH: Henry’s constant.

### 2.2. Environmental risk assessment

Risk assessment is based on an approach that combines two fundamental components of risk: the hazard, defined as the a.s. potential to cause adverse effects to non-target organisms; and the exposure, corresponding to the likelihood that these non-target organisms encounter the substance.

These two components are integrated into a composite indicator, designated as the Risk Index (RI; equation 1). The hazard component comprises the toxicity factor (T) and the bioaccumulation factor (B), while the exposure component is composed of the estimated quantity of a.s. (Q), the mobility factor (M), and the persistence factor (P) in the specific compartment.

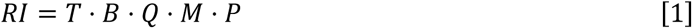

Each factor is estimated independently using models or empirical relationships, as described in Corder (2025b). To ensure reproducibility and applicability to Walloon agriculture purposes, methodological adaptations were introduced to focus on the most influential parameters for which reference or estimated values are available, despite the lack of detailed information on application conditions, transfer processes, or degradation pathways.

The model produces a relative risk index, that enables the comparison of each a.s. potential contribution to the overall risk across four environmental compartments characterized by specific transfer and dissipation processes:

- **Surface water compartment (SW)**, corresponding to visible freshwater bodies, flowing or stagnant (rivers, streams, lakes, canals), together with their associated aquatic ecosystems;
- Groundwater compartment (GW);
- **Non-cultivated terrestrial compartment (NT)** encompassing all terrestrial environments outside cultivated soils and providing potential habitats for biodiversity (mammals, birds, insects, soil fauna and flora);
- **Cultivated soil compartment (CS)**, referring to agricultural soils directly exposed to PPPs and associated with soil biota (focusing on earthworms).

As no natural ecosystem is associated with groundwaters, T and B factors are not determined for this compartment. The calculated index only estimates a compartment exposure to a.s. and is therefore named Exposure Index (EI), as defined in equation 2.

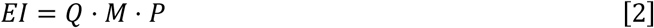

The M and P factors integrate pedological, hydrological, and meteorological conditions, as well as the a.s. application season and the crop to which they are applied.

#### 2.2.1. T factor: toxicity

The *T* factor evaluates the toxic potential of an a.s. toward non-target organisms inhabiting the ecosystems associated with each environmental compartment. According to Kooijman (1987) and Aldenberg et al. (1993), toxicity values (e.g. LC_50_, LD_50_, EC_50_) of an a.s. towards organisms of an ecosystem are expected to follow a log-normal distribution. Therefore, available toxicity data for each a.s. are aggregated accordingly for each environmental compartment considered.

Variables derived from acute toxicity tests are standardized at the species level by dividing each value by a reference toxicity threshold specific to the target species obtained from the PPDB. When multiple species within the same taxon are available, the mean of the standardized values is calculated to represent that taxon (*Tj*). The mean taxon-specific values are subsequently transformed using an inverse logarithmic function. Finally, the arithmetic mean of the transformed values of all representative taxa of a given environmental compartment is calculated.

Given the uncertainty arising from data variability and the presence of missing values, the *T* factor needs to be adjusted upward following the degree of uncertainty. The standard deviation associated with the logarithmic mean is used to calculate an uncertainty coefficient (ε). This coefficient is derived from Student’s *t*-distribution, applied to a sample of *n* taxa representative of the ecosystem, with an 80% confidence level.

The *T* factor for an ecosystem comprising *n* taxa is calculated using equation 3.

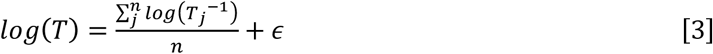

This method is applied to the following taxa, depending on the environmental compartment:

- **SW compartment**: fishes, aquatic invertebrates, crustaceans, aquatic plants, sediment-dwelling organisms, and algae;
- **NT compartment**: bees, earthworms, birds, and mammals;
- **CS compartment**: earthworms.

For the CS compartment, only tests conducted on earthworms are available for a sufficient number of a.s. Because the calculation is based on only one taxon, the computation of an upper bound and a confidence interval is not applicable. On the other hand, the T factor is not evaluated for the GW compartment, as it is not associated to a natural ecosystem.

#### 2.2.2. B factor: bioaccumulation

At equivalent toxicity and exposure, a highly bioaccumulative substance is more hazardous to ecosystems (van der Werf, 1997), as it may induce delayed or persistent effects, even at low environmental concentrations.

The bioaccumulation factor (B) integrates bioconcentration, dietary bioaccumulation and biomagnification along the trophic chain. This factor represents the potential of an a.s. to accumulate in living organisms. Using quantitative structure–activity relationship (QSAR) models, a.s. bioaccumulation is estimated for a high trophic-level organism representative of each environmental compartment, based on a.s. physicochemical properties like octanol-water (K_OW_) and octanol-air (K_OA_) partition coefficients:

- **SW compartment**: a biomagnification model for the rainbow trout, a representative high trophic-level fish species, is used (Arnot et al., 2003);
- **NT compartment**: a biomagnification model for the Arctic wolf, representative of a high trophic-level terrestrial species, is used (Gobas et al., 2003);
- **CS compartment**: a bioconcentration model for earthworms derived from Pflugmacher (1992) is used. This model has been adapted to prevent overestimation of bioaccumulation for highly hydrophobic substances (log *Kow > 6*) in the original model.

These QSAR models estimate a.s. bioaccumulation coefficients (C_b_), later used in the calculation of the *B* factor for the different environmental compartments following equation 4.

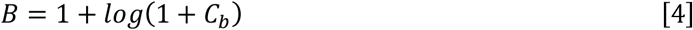

The absence of bioaccumulation (i.e., when *Cb = 0*), the *B* factor retains a minimum value of 1. Bioaccumulation is hence considered an aggravating factor to the risk.

#### 2.2.3. Q factor: quantity used

The *Q* factor represents the quantity of a.s. considered in the risk assessment. Two approaches to estimate this factor are distinguished depending on the objective of the analysis:

i. Temporal risk assessment. In this case, *Q* corresponds to the total annual quantities of substances used (kg·yr⁻¹) across the 15 agricultural sectors associated with arable crops.
ii. Prospective comparison of substances under different scenarios. In this framework, the *Q* factor is defined as a reference application dose (kg·ha⁻¹·yr⁻¹). This dose corresponds to the maximum authorized dose per application and per hectare for a given crop and, where relevant, for a specific target.

#### 2.2.4. M factor: mobility

The *M* factor assesses the probability of an a.s. to be transferred to SW and GW compartments, as well as the resulting exposure of organisms associated with CS and NT environments. Its expression is compartment-specific in the model, depending on the dominant transfer processes (runoff and erosion, leaching, spray drift, or volatilization) (**Figure 1**). For the non-cultivated terrestrial compartment, the mobility of organisms toward treated areas is also considered.

**Figure 1:**
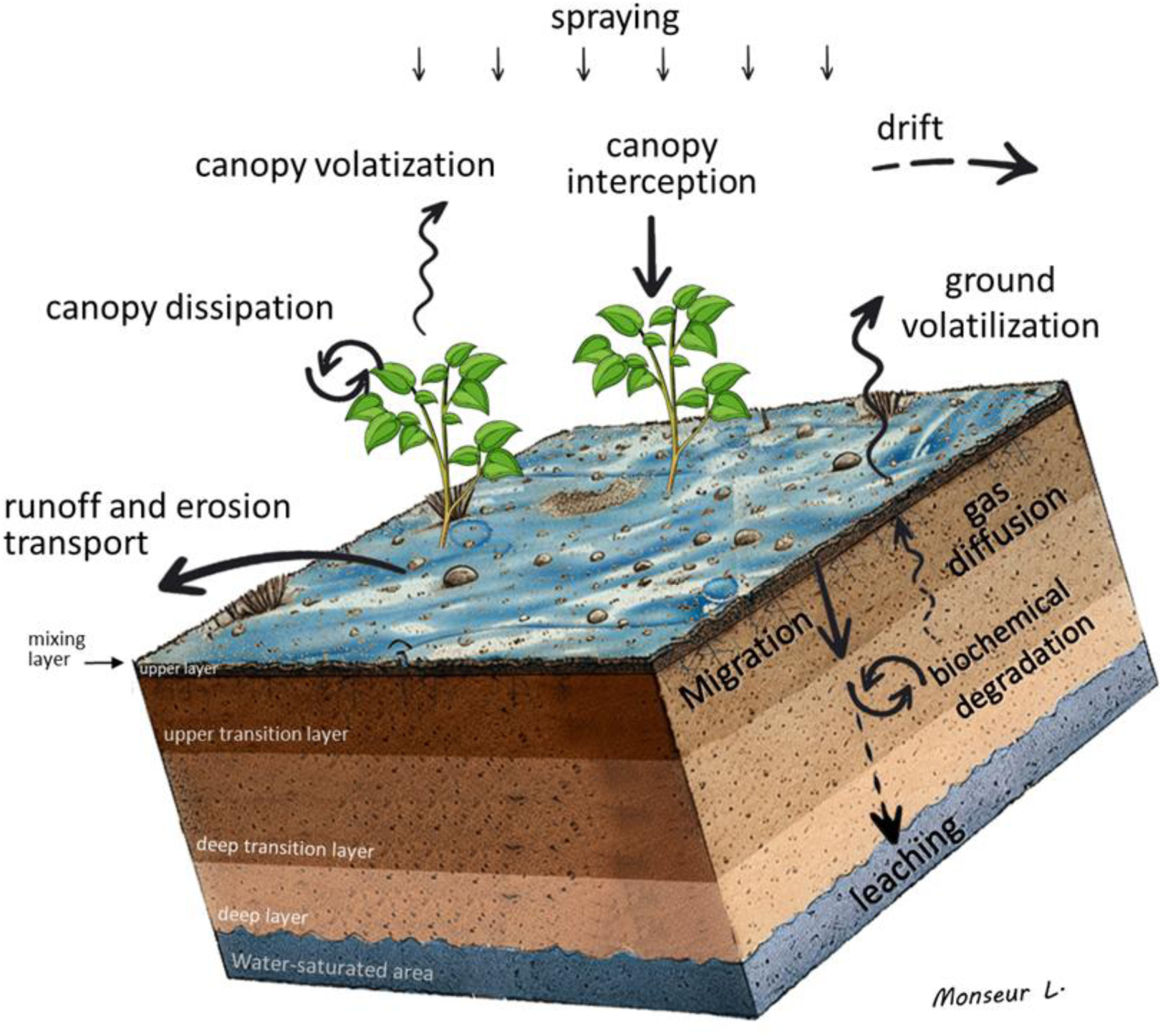
Transfer and dissipation processes of active substances applied in arable cropping systems considered in the Phytorisque. Following spraying, pesticides may be intercepted by the plant canopy, volatilize, drift, or reach the soil surface. In the soil, they may undergo degradation, gas diffusion, and vertical migration through soil layers, potentially leading to leaching toward groundwater. Surface transport may also occur through runoff and erosion, contributing to off-site contamination.

Transfer processes were modeled integrating soil hydropedological conditions (notably soil texture, slope, and organic matter content), a.s. physicochemical properties, meteorological conditions prevailing during the year of application. The season of application is also considered as it affects weather conditions and the crop’s developmental stage at the time of application. The values of the M factors presented in this study are derived from arable cropping systems, aggregated across the 15 analyzed sectors. To obtain a representative value at the scale of all arable crops, a weighted average of the sector-specific values was calculated based on the quantities of a.s. estimated for each sector in Corder (2025a). Furthermore, application periods of the a.s. was collected from sector experts of the 15 representative arable crops, and a frequency-weighted mean of the seasonal results was used to derive the final factor values (Corder ASBL, 2025b).

For the SW compartment, the Msw factor quantifies the fraction of the applied a.s. exported from the application area to surface water bodies via runoff, in both dissolved and particulate forms. This fraction is defined as the product of the quantity of a.s. not intercepted by the crop canopy and the runoff transfer potential (*Fsw*). Runoff is therefore the only transport pathway towards SW taken into consideration in Phytorisque. Although other transport pathways (e.g., drainage, drift, and volatilisation) exist, they have been estimated negligible by comparison with runoff for the indicator developed in this study.

To calculate this transfer potential, runoff is estimated using the Curve Number method (McCuen, 1983; Forootan, 2023), and erosion is assessed using the Modified Universal Soil Loss Equation (Kruijne et al., 2011). The modeling of substance transfer is further performed as described in Leonard (1990), Huber et al. (1998) and Young et al. (2019). The M_SW_ factor is calculated according to equation 5.

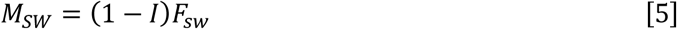

where *I* is the interception factor of the substance by the crop canopy, which depends on the crop’s developmental stage and therefore the application season, and *F_SW_* is the runoff transfer potential.

For the GW compartment, the *M_GW_* factor assesses the fraction of a.s. not intercepted by the crop canopy and leached beyond a depth of two meters. Beyond this threshold, the substance is considered to have exited the biologically active soil zone, where biological degradation has a significant effect.

The transfer potential (*F_GW_*) is based on the attenuation factor proposed by (Hornsby et al., 1985), who modelled the soil as a multilayer profile. The detailed calculation procedure for this potential is presented in Corder (2025b).

This factor is therefore expressed according to equation 6.

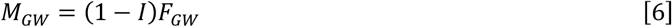

For the NT compartment, the *M_NT_* factor evaluates both a.s. dispersion within the environment and the movements of terrestrial organisms from their habitats toward treated areas. This factor is defined according to equation 7.

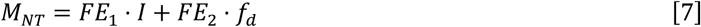

The first term of equation 7 accounts for the organisms’ exposure due to their movements in treated crops. The ecological exposure factor FE₁ represents the probability of non-target organisms’ presence in the treated area. It modulates the interception (I) depending on the ecological potential of the surrounding environment and its spatial proximity to arable crops. The second term of equation 7 assesses exposure resulting from the off-site dispersion of a.s. applied to crops. The dispersed fraction (*f_d_*) expresses the proportion of a.s. transferred to the environment through spray drift, volatilization, and erosion, while the ecological exposure factor *FE*₂ evaluates the relative ecological potential of the receiving environments surrounding the arable crops. The calculation method for the exposure factors (*FE*₁ and *FE*₂) and the dispersed fraction (f_d_), are detailed in Corder (2025b).

For the CS compartment, the *M_CS_* factor corresponds to the a.s. fraction reaching the soil during application. This factor is therefore calculated using equation 8.

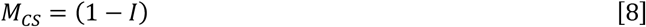

#### 2.2.5. P factor: persistence

The *P* factor evaluates the average persistence of an a.s. in an environmental compartment prior to its dissipation. This duration influences the probability of contact between the a.s. and the non-target organisms in that compartment.

Persistence is modelled using a kinetic function describing the evolution of the a.s. residual fraction with time in the compartment after its initial transfer. The evolution of the non-dissipated fraction of an a.s is expressed by a generic function *f(t)* defined in equation 9.

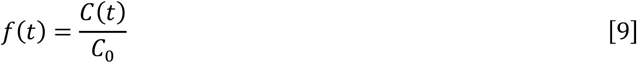

where *C(t)* is the a.s. concentration at time *t*, and *C*₀ is the initial concentration of a.s. The *P* factor is calculated as the statistical expectation of this dissipation function, i.e., the first-order moment of the function *f*′(*t*), according to equation 10.’

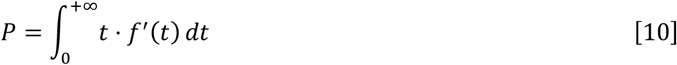

Different kinetic models are applied depending on the environmental compartment and the dominant dissipation processes. These models are described in detail in Corder ASBL *et al*. (2025b).

The season of application intervenes to adjust transfer and dissipation processes, as these are influenced by temperature, precipitation, solar radiation, and soil moisture content. The P factors used in this study are derived from arable cropping systems, aggregated across the 15 analyzed sectors. The representative value at the scale of all arable crops was calculated based on the weighted average of a.s used in a specific sector as described in Corder (2025a).

For the SW compartment, in the absence of sufficient data on the a.s. biodegradation kinetics in aquatic environments, persistence is estimated based on abiotic processes, namely hydrolysis and photolysis. The concentration is assumed to follow first-order exponential decay kinetics, with an apparent constant rate corresponding to the weighted sum of the rate constants of these two processes. Hydrolysis rate is adjusted according to the mean seasonal temperature of river water using the Arrhenius equation, as suggested by EFSA (2008). Photolysis rate is expressed as a function of solar irradiance, which depends on latitude and time of year, and decreases with increasing water depth (Zepp et al., 1977). A representative mean depth for surface waters in Wallonia was therefore used to adjust the attenuation of photolysis rate.

Finally, an additional constant dissipation rate is incorporated into the model to account for hydrological renewal induced by flow dynamics and biodegradation, thereby limiting persistence to a seasonal timescale.

For the GW compartment, degradation occurs through hydrolysis. The model is based on first-order kinetics, and the degradation rate constant is adjusted according to the mean groundwater temperature, in accordance with the Arrhenius equation, as recommended by EFSA (2008).

For the NT compartment, persistence reflects the heterogeneous structure of this compartment, comprising two sub-compartments: (i) the crop canopy, where part of the applied quantity is intercepted, and (ii) the adjacent non-treated area to which part of the remaining applied quantity is transferred through erosion from the treated area. Overall dissipation is modelled using a DFOP (Double First-Order in Parallel) kinetic approach, combining two dissipation rates occurring in the two sub-compartments (Boesten et al., 2006). Due to the limited availability and lack of harmonized data on pesticide dissipation from crop canopies, a simplified parameterization was adopted. For highly volatile substances, canopy dissipation is assumed to be mainly driven by volatilization. For less volatile substances, a median dissipation rate derived from PPDB data is applied (k = 0.15 day⁻¹). This dissipation rate was determined following an analysis of foliar residue half-lives reported in the PPDB (n = 180), which indicated that most a.s. persist on foliage for 2–14 days (median foliar residue DT₅₀ = 6.7 days).

For the CS compartment, persistence is modelled using variable-rate kinetics applied to a multilayer system integrating three dissipation processes: (i) leaching of the a.s., (ii) its biochemical degradation in soil, and (iii) its volatilization into the atmosphere.

In the model, biochemical degradation is assumed to be constant within the upper 10 cm of the soil and to decrease exponentially below this depth. This assumption is supported by the observed correlation between organic matter content and biological activity in the soil, together with the exponential decline of organic matter with increasing soil depth (Wilson et al., 1993; Murphy et al., 1998).

Each soil horizon is characterized by specific physicochemical conditions, such as organic matter content, temperature, moisture, biological activity intensity, and gas diffusion time to the surface. Consequently, leaching, biochemical degradation, and volatilization are modulated by the average depth reached by the a.s. at time t. The overall dissipation rate depends on the average position of the substance in the soil profile. Since analytical integration of equation 10 is not possible in this case, the solution is obtained through numerical integration, discretized into daily steps.

#### 2.2.6. Phytorisque indices

By combining the five factors described above, the Phytorisque model can be used to calculate three types of environmental risk indices, each suited to different purposes:

1. **Monitoring indices** are linked to the Aggregated Risk Index (ARI), obtained by aggregating the RI calculated for each a.s. They track changes in environmental risk over time for each compartment at the regional level, associated with the use of PPP in a specific agricultural sector or in a group of sectors. Normalized to a reference year, they evaluate changes in the global risk of PPP use over time.
2. **Diagnostic indices** identify the factors contributing to risk and assess the relative contribution of each a.s. to the overall risk. This contribution is calculated by relating the RI of an a.s. (equation 1) to the ARI. It is particularly relevant for identifying, for a given year and for each environmental compartment, the substances that contribute most to the overall risk.
3. **Prospective assessment indices** are associated with the Potential Risk Index (PRI), in which the quantity used (*Q* factor) corresponds to the reference dose (expressed in kg·ha⁻¹·yr⁻¹) of the a.s. (see Section 2.3). This index allows comparing the level of risk associated with a standardized dose, regardless of the quantities used at the regional level. It is particularly relevant for anticipating substance substitution mechanisms or for analyzing specific crop management scenarios. The PRI is a relative index that can be expressed as an absolute value or normalized to a reference substance.

### 2.3. Sensitivity analysis of the Phytorisque transfer models

A sensitivity analysis of Phytorisque transfer model was performed against a.s. physicochemical properties. Sensitivity to physicochemical parameters (K_OC_ and DT_50_) of a.s. was evaluated using a univariate approach, in which one parameter was varied at a time while the others were kept at median values, defined from the distribution of substances used in arable crops. In this analysis, meteorological conditions from 2023 were used, as this year is considered representative of average climatic conditions in Wallonia. The most relevant findings are reported in the results section, and more detailed results are presented in Corder (2025b).

## 3. RESULTS

### 3.1. Hydrological, pedological and meteorological context of Wallonia

The main hydropedological and climatic parameters used in the Phytorisque model are presented in Table 2.

**Table 2:** Pedological, hydrological, and meteorological parameters used for the calibration of the Phytorisque model, representative of arable cropping systems in Wallonia (Year 2022).

| Parameter | Value | Statistic |
| --- | --- | --- |
| *Hydropedological parameters representative of the main crops* |  |  |
| Soil water content at field capacity [-] | 0,37 | Median |
| Slope [%] | 3,5 | Median |
| Organic carbon content (0–10 cm) [%] | 1,8 | Mean |
| dry soil density [$g \cdot cm^{-3}$] | 1,33 | Mean |
| Soil erodibility factor [$t \cdot h \cdot mm^{-1} \cdot MJ^{-1}$] | 0,06 | Median |
| *Climatic parameters (station of Sombrefe, 2022)* |  |  |
| Annual precipitation [mm] | 496 | - |
| Mean precipitation return period [days] | 6,0 | Mean |
| Mean annual temperature [$^{\circ}C$] | 11,6 | Mean |

### 3.2. Risk assessment for surface water

The hazard factors (toxicity and bioaccumulation) and exposure factors (mobility and persistence) were calculated in the SW compartment for the five selected a.s. used on arable crops in Wallonia together with the RI (Table 3). Among the tested a.s., lambda-cyhalothrin has the highest toxicity and bioaccumulation values, and chlorantraniliprole and maleic hydrazide have high mobility and persistence values. Moreover, lambda-cyhalothrin has the highest RI, while maleic hydrazide remains with the lowest value.

**Table 3:** Toxicity (T), Bioaccumulation (B), Mobility (M), Persistance (P) and Risk Index for surface waters (RIsw) as obtained from the Phytorisque model. M and P for each active substance are aggregated values derived from 15 representative arable crops grown in Wallonia. Calculations were performed using meteorological conditions for 2022 in Wallonia, regional hydropedological conditions, and estimated quantities of PPP applied to 15 arable crops in 2022 in Wallonia.

| Active substance | T [-] | B [-] | M [%] | P [days] | R<sub>sw</sub> |
| --- | --- | --- | --- | --- | --- |
| Bromuconazole | 42,9 | 3,7 | 0,35 | 50 | $5,53 \times 10^4$ |
| Chlorantraniliprole | $1,12 \times 10^3$ | 3,2 | 0,71 | 10 | $8,83 \times 10^4$ |
| Clomazone | 33,7 | 2,9 | 0,40 | 77 | $9,65 \times 10^4$ |
| Maleic hydrazide | 0,3 | 1,3 | 0,08 | 79 | $6,01 \times 10^2$ |
| Lambda-cyhalothrin | $9,93 \times 10^5$ | 7,2 | 0,27 | 74 | $6,87 \times 10^8$ |

The transfer potentials to surface waters (F_sw_) for the five a.s. are presented in Figure 2. Transfers were estimated under two contrasting climatic conditions: the wet year 2021 and the particularly dry year 2022, to evidence the influence of meteorological conditions on pesticide transport. Overall, transfer potentials were substantially higher in 2021 than in 2022 for all a.s., reflecting the strong influence of rainfall on runoff and erosion processes. The relative contribution of each transport pathway, runoff or erosion, differed between a.s.; lambda-cyhalothrin was predominantly transported in particulate form through erosion processes, whereas maleic hydrazide was mainly transported in dissolved form via runoff.

**Figure 2:**
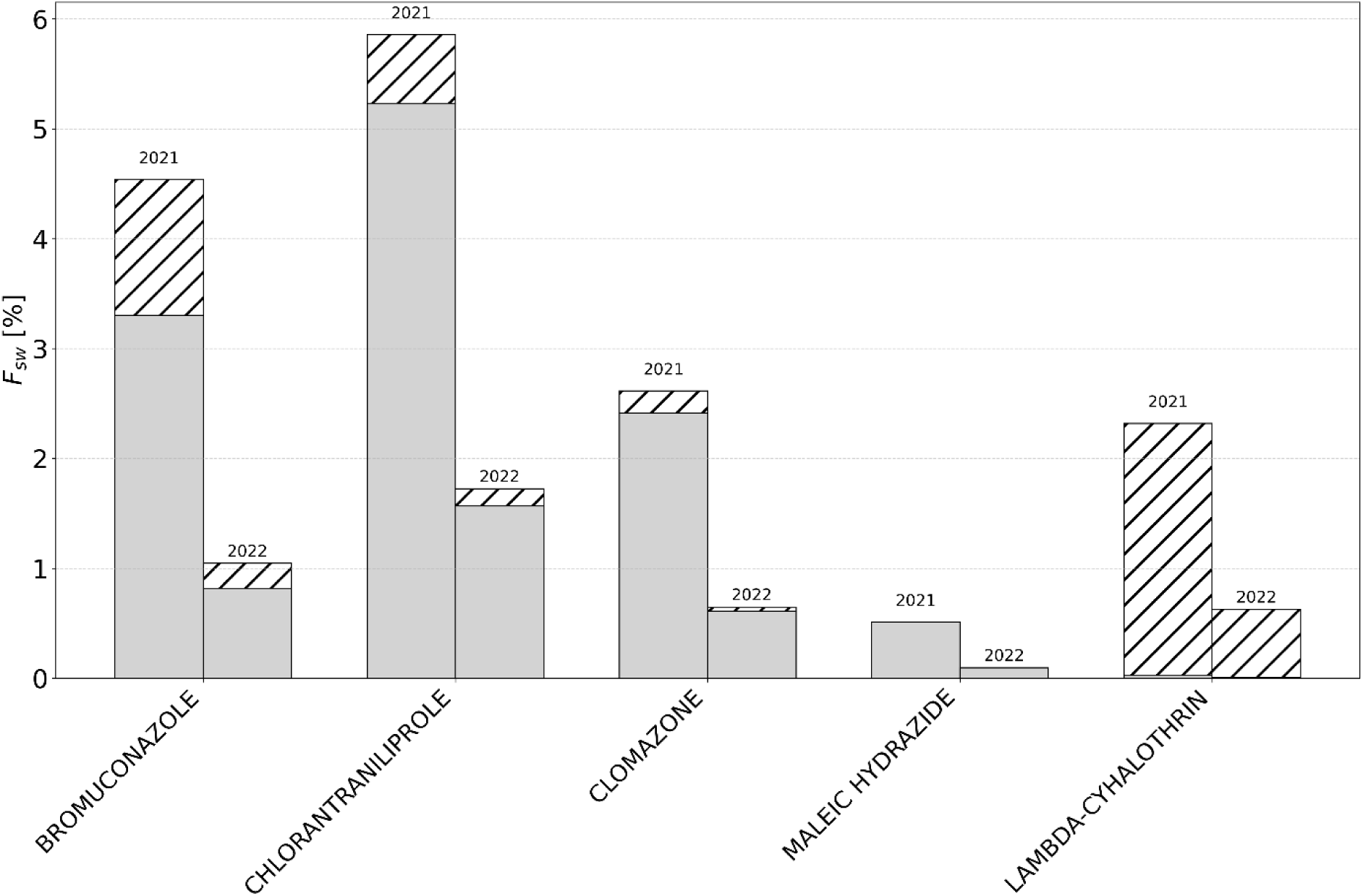
Transfer potential to surface waters (FSW) for five a.s. as assessed by the Phytorisque model under contrasting climatic conditions (wet year 2021 and dry year 2022). Bars show the total transfer potential, split between dissolved transport via surface runoff (solid) and particulate transport associated with soil erosion (hatched). Calculations were performed using regional hydropedological conditions and estimated quantities of PPP applied to 15 arable crops in 2022 in Wallonia.

A sensitivity analysis of transfer potential to SW was performed with respect to soil persistence (soil DT₅₀) and sorption capacity (K_OC_). Figure 3 shows the transfer potential variation with soil DT_50_ at fixed K_OC_ values. The effect of soil DT_50_ is particularly visible at low soil DT_50_ values, and a plateau is observed as this parameter increases. With lower K_OC_ values, the plateau is observed at lower soil DT_50_.

**Figure 3:**
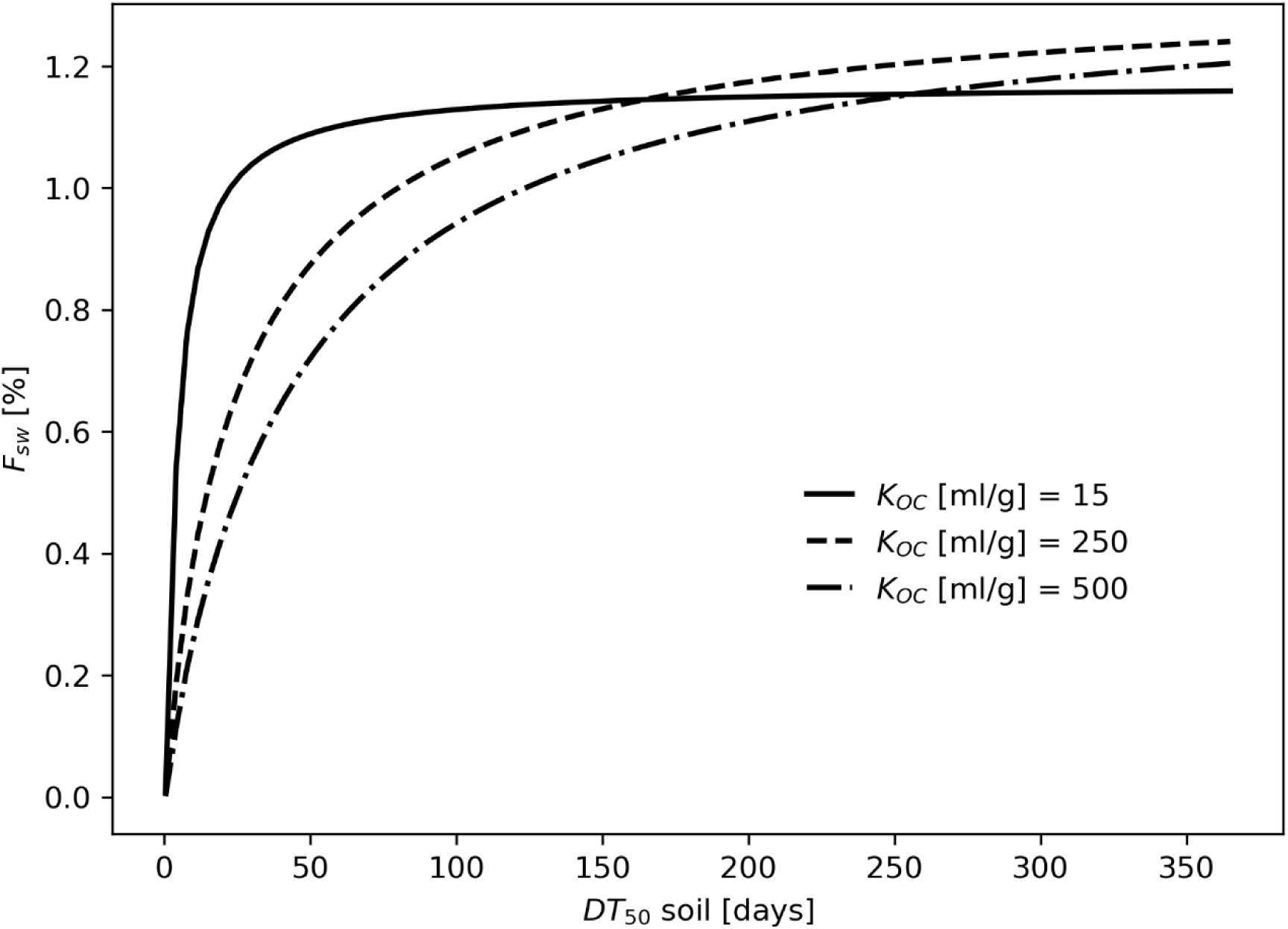
Sensitivity analysis of the transfer potential to surface waters (FSW) in the Phytorisque model to soil DT50 for three levels of soil adsorption coefficients (Koc = 15, 250, and 500 mL/g). Based on meteorological conditions of 2023 and hydropedological conditions of Wallonia.

Transfer potential sensitivity to K_OC_ was also investigated while fixing soil DT_50_ at 365 days (Figure 4). The transfer potential through the dissolved and particulate phases are distinguished. The dissolved phase transfer potential decreases with increasing K_OC_, whereas particulate phase transfer potential increases. Consequently, the dominant transport pathway shifts from dissolved transport at low K_OC_ values to particulate transport at higher K_OC_ values.

**Figure 4:**
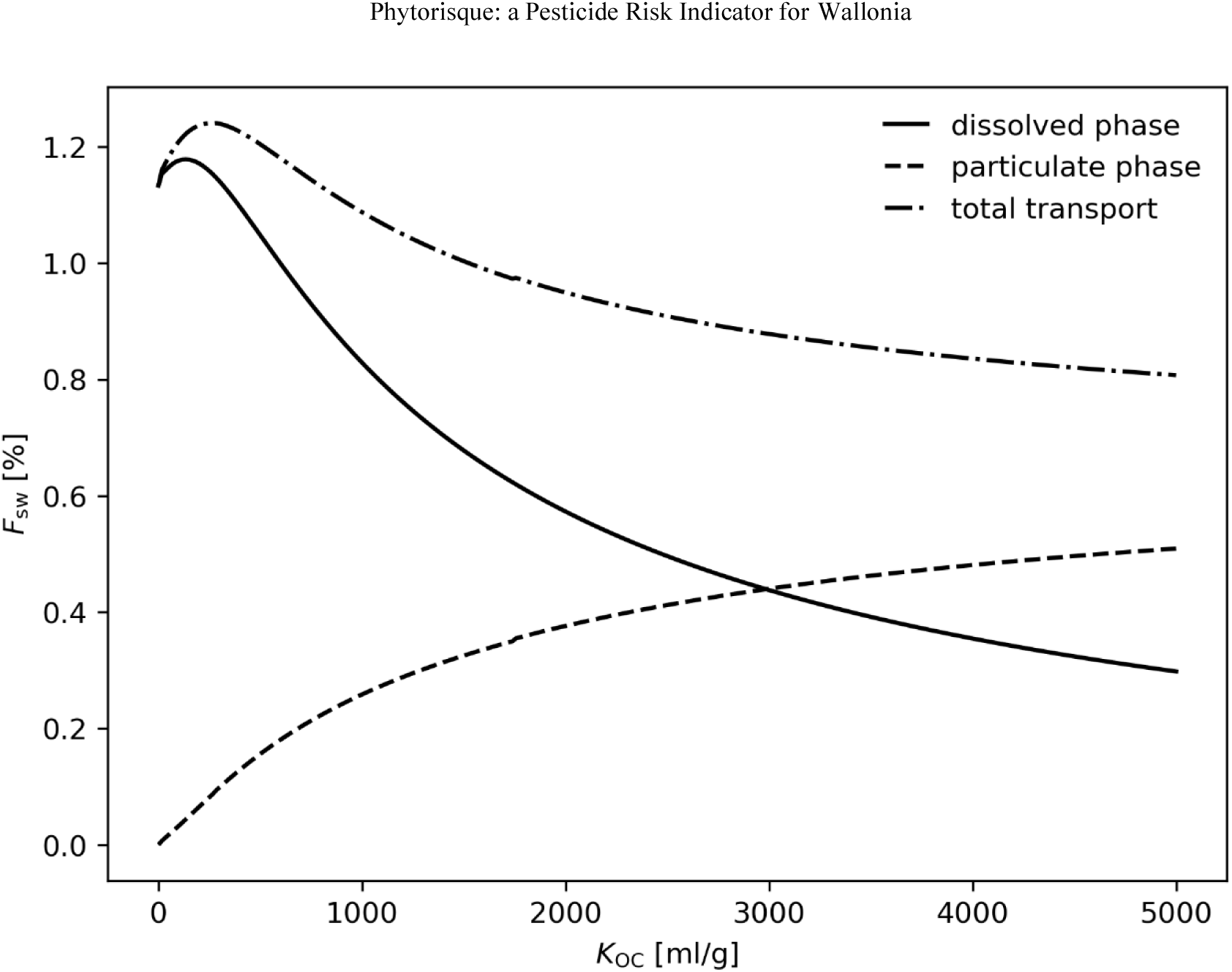
Sensitivity of the transfer potential to surface waters (FSW) in the Phytorisque model to the soil adsorption coefficient (KOC) (soil DT₅₀ = 365 days). The total transfer potential (dash-dot line) reflects contributions from dissolved transport via runoff (solid line) and particulate transport via soil erosion (dashed line). Calculations were based on 2023 meteorological conditions and regional hydropedological conditions in Wallonia.

### 3.3. Groundwater exposure assessment

As GW is not associated with a natural ecosystem, hazard factors are not calculated for this compartment and only exposure-related parameters are considered. The exposure factors (mobility and persistence), together with the Exposure Index (EI) for the five a.s., are presented in Table 4. Among the selected a.s., chlorantraniliprole exhibits both high mobility and persistence, resulting in the highest EI. In contrast, lambda-cyhalothrin shows negligible mobility and consequently an EI equal to zero (Table 4).

**Table 4:** Mobility (M), Persistance (P) and Exposure Index for groundwaters (EIGw) as obtained from the Phytorisque model. M and P for each active substance are aggregated values derived from 15 representative arable crops grown in Wallonia. Calculations were performed using meteorological conditions for 2022 in Wallonia, regional hydropedological conditions, and estimated quantities of PPP applied to 15 arable crops in 2022 in Wallonia.

| Active substance | M [%] | P | $IE_{GW}$ |
| --- | --- | --- | --- |
|  |  | [days] |  |
| Bromuconazole | $1,60 \times 10^{-3}$ | 101 | $3,09 \times 10^2$ |
| Chlorantraniliprole | $3,88 \times 10^{-2}$ | 1 233 | $1,51 \times 10^4$ |
| Clomazone | $1,1 \times 10^{-10}$ | 1 233 | $4,27 \times 10^{-4}$ |
| Maleic hydrazide | $6,25 \times 10^{-45}$ | 1 233 | $1,95 \times 10^{-37}$ |
| Lambda-cyhalothrin | 0 | 567 | 0 |

In addition, Figure 5 shows a.s. transfer potential sensitivity to GW with K_OC_ and soil DT_50_. Transfer potential decreased drastically with increasing K_OC_. Poorly retained a.s. (low K_OC_ values) have the highest transfer potential, particularly when persistence in soil is high. In contrast, the transfer potential becomes negligible for highly sorbing substances (K_OC_ > ∼300–400 mL g⁻¹). Increasing soil DT₅₀ leads to substantially higher transfer potentials, with the highest values observed for soil DT_50_ = 365 days.

**Figure 5.**
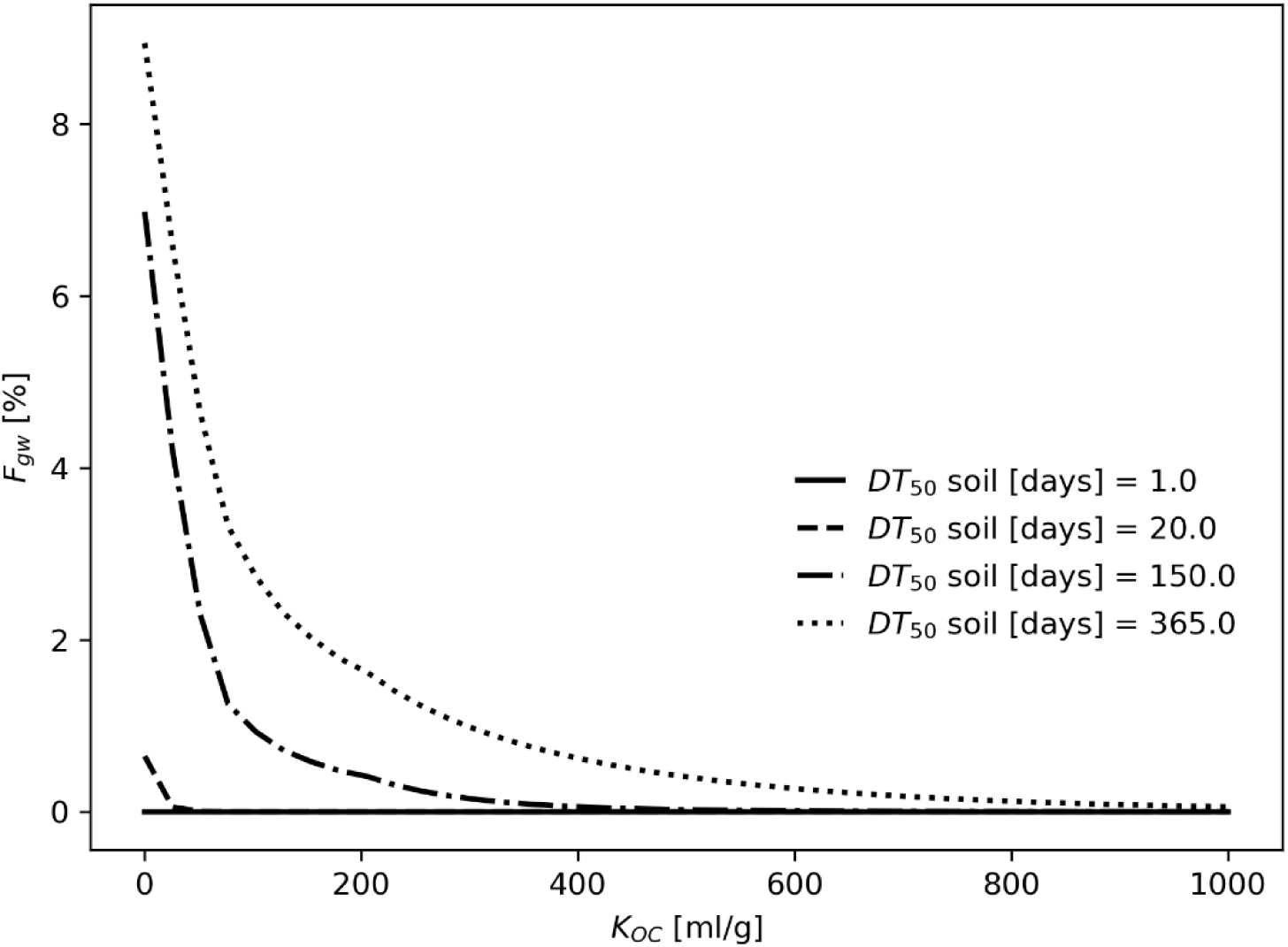
Sensitivity of the transfer potential to groundwaters (FGW) in the Phytorisque model to the soil adsorption coefficient (KOC) (soil DT₅₀ fixed at 1, 20, 150 and 365 days). Calculations were based on 2023 meteorological conditions and regional hydropedological conditions in Wallonia.

### 3.4. Risk assessment for non-cultivated terrestrial compartment

Values for the hazard factors, the exposure factors and the RI in the NT compartment are presented in Table 5. Lambda-cyhalothrin has the highest toxicity and bioaccumulation potentials, resulting in the highest RI among the tested a.s. In contrast, maleic hydrazide shows low toxicity, bioaccumulation, mobility and persistence. Among the selected a.s., clomazone has the lowest RI.

**Table 5:** Toxicity (T), Bioaccumulation (B), Mobility (M), Persistance (P) and Risk Index for the non-cultivated terrestrial compartment (RI_NT_) as obtained from the Phytorisque model. M and P for each active substance are aggregated values derived from 15 representative arable crops grown in Wallonia. Calculations were performed using meteorological conditions for 2022 in Wallonia, regional hydropedological conditions, and estimated quantities of PPP applied to 15 arable crops in 2022 in Wallonia.

| Active substance | T [-] | B [-] | M [-] | P [days] | $RI_{NT}$ |
| --- | --- | --- | --- | --- | --- |
| Bromuconazole | 2,7 | 2,6 | 1,07 | 143 | $6,31 \times 10^5$ |
| Chlorantraniliprole | 2,6 | 2,3 | 0,94 | 37 | $3,63 \times 10^4$ |
| Clomazone | 4,1 | 1,5 | 1,17 | 6 | $3,17 \times 10^4$ |
| Maleic hydrazide | 1,0 | 1,0 | 0,55 | 6 | $9,21 \times 10^4$ |
| Lambda-cyhalothrin | 82,1 | 3,1 | 1,13 | 95 | $2,87 \times 10^6$ |

The M_NT_ factor integrates two exposure pathways for terrestrial organisms: direct exposure when terrestrial organisms move across treated crops and indirect exposure resulting from the dispersion of a.s. outside the treated areas through processes such as drift, volatilization, and erosion. These two pathways are distinguished for the 5 selected a.s. in Figure 6. For most a.s., exposure is dominated by direct contact within treated areas. Among them, Clomazone is the most subjected to dispersion in the environment, resulting in a nearly equal contribution between the two exposure pathways.

**Figure 6:**
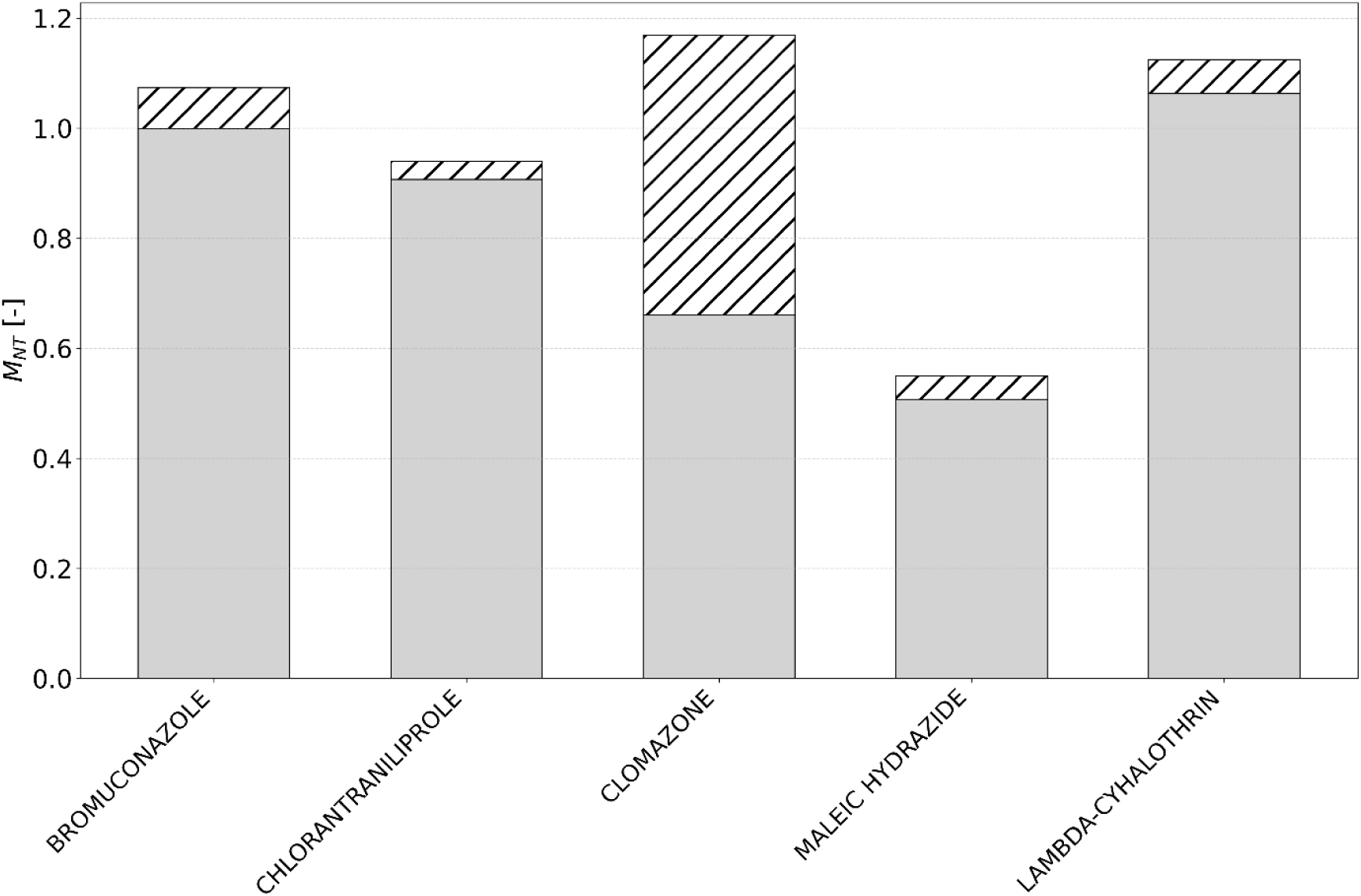
Mobility factor for the NT compartment (M_NT_) of the five selected a.s. as assessed by the Phytorisque model. M_NT_ can be split between exposure resulting from the movement of terrestrial organisms across treated crops (solid bars) and exposure due to a.s. dispersion outside the treated area (hatched bar). Calculations were performed using meteorological conditions for 2022 in Wallonia, regional hydropedological conditions, and estimated quantities of PPP applied to 15 arable crops in 2022 in Wallonia.

### 3.5. Risk assessment for cultivated soil compartment

The hazard and exposure factors, together with the resulting RI, for the five selected a.s. in the CS compartment are presented in Table 6. Bromuconazole is the most persistent in soil and has consequently the highest RI for this compartment. Maleic hydrazide has the lowest persistence and RI values.

**Table 6:** Toxicity (T), Bioaccumulation (B), Mobility (M), Persistance (P) and Risk Index for cultivated soils (RICS) as obtained from the Phytorisque model. M and P for each active substance are aggregated values derived from 15 representative arable crops grown in Wallonia. Calculations were performed using meteorological conditions for 2022 in Wallonia, regional hydropedological conditions, and estimated quantities of PPP applied to 15 arable crops in 2022 in Wallonia.

| Active substance | T [-] | B [-] | M [-] | P [days] | RI<sub>CS</sub> |
| --- | --- | --- | --- | --- | --- |
| Bromuconazole | 2,0 | 2,3 | 0,40 | 1772 | 6,22 x 10<sup>6</sup> |
| Chlorantraniliprole | 1,0 | 1,9 | 0,46 | 981 | 2,71 x 10<sup>5</sup> |
| Clomazone | 12,8 | 1,7 | 0,60 | 52 | 2,12 x 10<sup>6</sup> |
| Maleic hydrazide | 1,0 | 1,0 | 0,70 | 4 | 8,09 x 10<sup>4</sup> |
| Lambda-cyhalothrin | 2,0 | 4,7 | 0,36 | 485 | 8,10 x 10<sup>5</sup> |

P_CS_ sensitivity with respect to K_OC_ and soil DT_50_ is presented in Figure 7. For all K_OC_ values, P_CS_ increases with soil DT₅₀. A plateau is reached upon soil DT_50_ increase. The value of this plateau depends on K_OC_. Overall, at high soil DT_50_, variations in persistence of the a.s. are mainly driven by K_OC._

**Figure 7.**
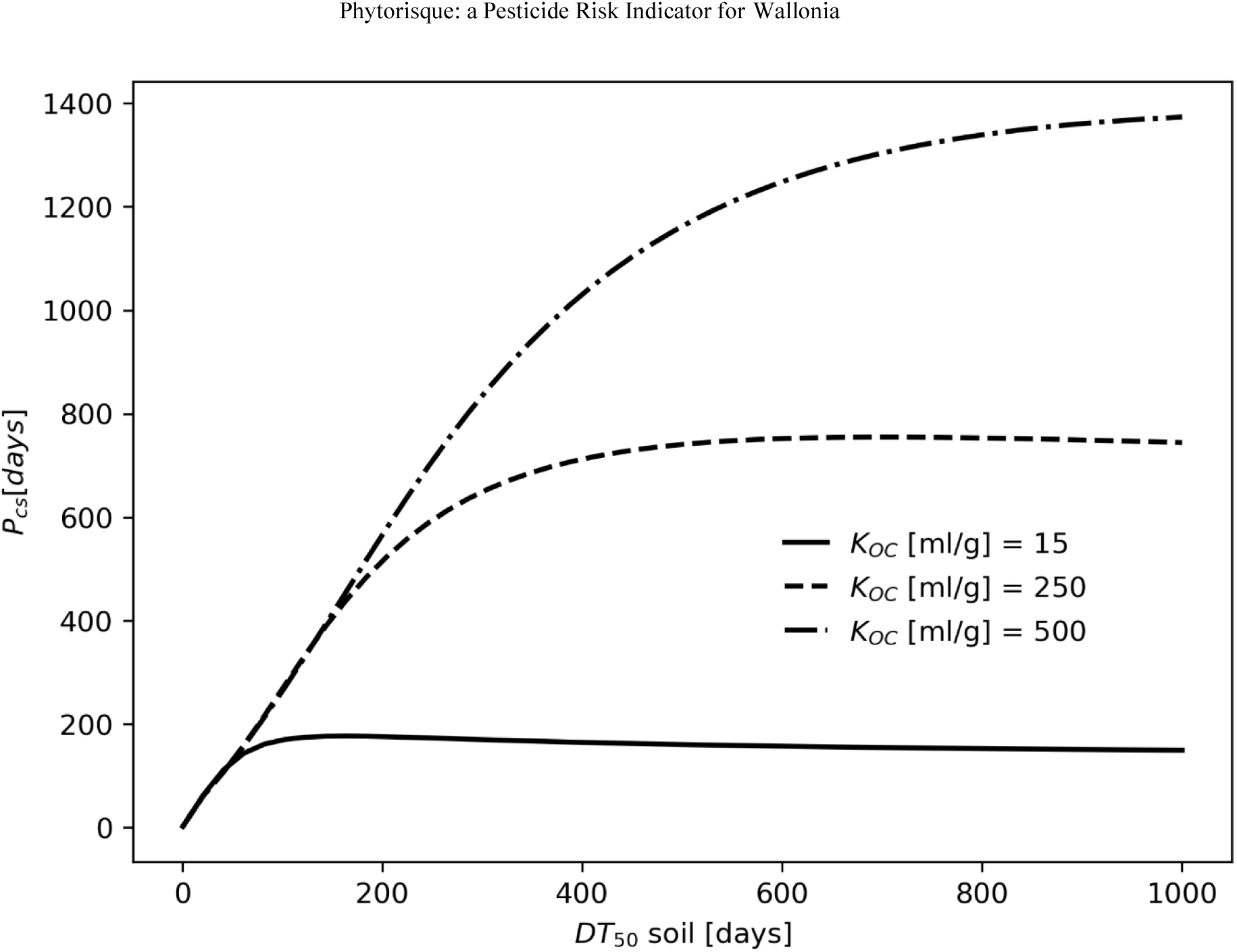
Sensitivity of the Persistence factor in cultivated soils (P_CS_) to soil DT_50_ for different soil adsorption coefficient (K_OC_ = 15, 250 and 500). Calculations were based on 2023 meteorological conditions and regional hydropedological conditions in Wallonia.

## 4. DISCUSSION

### 4.1. Conceptual specificity of Phytorisque

The Phytorisque developed in this study follows an ETR-based approach to evaluate the risk associated with the use of PPP in agriculture. Traditionally, ETR expresses risk as the ratio between an exposure level and a toxicity threshold. In the present case, this logic is reformulated in a multiplicative framework (equation 1), where the RI is defined as the product of several factors describing both exposure and s.a. hazard effects. In the Phytorisque, the Q, M and P factors characterize the environmental exposure potential, whereas T and B factors reflect the intrinsic sensitivity of organisms and the potential for accumulation in non-target organisms. In particular, the T factor is defined as the inverse of toxicological values (equation 2), such that lower ecotoxicological thresholds correspond to higher T values. Accordingly, despite its multiplicative formulation, the Phytorisque can be interpreted as an ETR-based risk indicator.

Nowadays, most ETR-based indicators as well as methodologies used for homologation purpose are based on the estimation of a.s. predicted environmental concentration and non-effect concentration (Damalas et al., 2011; Strassemeyer et al., 2017; van Straalen et al., 2022). Instead, the Phytorisque model evaluates the load of a.s. transferred to each environmental compartment. Unlike previous indicators, it considers the cumulative effects of fluxes and dissipation processes, rather than determines concentrations at a specific time in each compartment. This methodological approach is justified by two main considerations:

i. First, considering the a.s. concentration at a regional level remains challenging, as converting transferred mass into environmental concentration depends on poorly constrained environmental parameters (Aravinna et al., 2006). Considering the most probable conditions in terms of PPP application, hydropedological and climatic conditions, the Phytorisque estimates the probability of transfer towards each environmental compartment considered. It could be interpreted as a proxy for off-site transfer and exposure potential rather than as a direct estimate of environmental concentrations.
ii. Second, instantaneous concentrations of a.s. cannot represent the cumulative and prolonged nature of transfer processes (Boesten et al., 2007). Two scenarios involving the same quantity of a.s. transferred to the environment may result in different exposure profiles, characterized by either short-term peak concentrations or diffuse and prolonged transfer. The Phytorisque model accounts for both processes, as illustrated in Figure 4. For example, a.s. with a strong affinity for soil particles are poorly mobilised by runoff and their transfer toward surface waters during the first rainfall event following application is limited. However, when such substances are also persistent, they remain in the soil and are available for particulate erosion over extended periods beyond the initial post-application event. Consequently, peak concentrations in surface waters remain negligible, but the cumulative mass flux transferred to aquatic ecosystems can be significant. This behavior is illustrated in figure 2 by lambda-cyhalothrin, which combines high K_OC_ with strong persistence. Therefore, this substance is more subject to prolonged, low-intensity transfer towards SW compartment.

Another important innovation of the Phytorisque lies in the explicit integration of bioaccumulation and persistence as aggravating factors in the risk assessment. The bioaccumulation factor, bounded below by 1, acts as an amplifying term that increases the estimated intrinsic hazard of a.s. by accounting for their potential to concentrate within biological organisms. Persistence, on the other hand, evaluates a.s. residence time in the environmental compartment considered, reflecting its sustained potential to cause adverse effects within that compartment. By incorporating these two factors, the Phytorisque provides a more comprehensive representation of long-term and cumulative risk dynamics.

### 4.2. Effects of physico-chemical properties on transfer processes

The influence of the physicochemical properties of a.s. on ecosystem exposure varies depending on the specific environmental compartment.

**For surface waters**, as reported by several authors (Leonard, 1990; Hornsby et al., 1993; Grill et al., 1999; Wendell et al., 2026), the sorption coefficient (K_OC_) and soil degradation (soil DT_50_) play a key role in the transfer of a.s. This transfer results from the combined contribution of dissolved and particulate transport pathways ((Commelin et al., 2022); Figures 3 and 4).

At low K_OC_ values, a.s. are weakly adsorbed to soil particles and are therefore prone to leaching toward groundwater (Reichenberger et al., 2007), notably in the loamy soils characteristic of Wallonia. This process competes with transfer toward surface waters, as a.s. are rapidly transported downward and become less available for interaction with surface runoff (Delvin et al., 2008; Silburn, 2023). At intermediate K_OC_ values, leaching becomes less significant and a larger fraction of the a.s. remains within the soil mixing layer. Interaction with surface runoff is enhanced, leading to a maximum transfer toward surface waters, as shown in Figure 4. At high K_OC_ values, strong sorption to soil particles limits transport in the dissolved phase. Transfer then occurs predominantly through erosion-driven particulate transport, whereby a.s. are carried by the soil particles to which they are adsorbed.

While dissolved transport mainly occurs during the first rainfall events following application, particulate transport is generally slower and becomes significant considering the contribution of rainfall events over a longer period. Consequently, a.s. characterized by both high K_OC_ and high soil DT_50_ are more likely to follow this pathway, as their persistence maintains their availability in shallow soil horizons for subsequent erosion events. Such observation was previously highlighted by several authors (Hornsby et al., 1993; Commelin et al., 2022).

**Regarding groundwater exposure**, K_OC_ and soil DT_50_ are among the dominant parameters influencing the leaching rate of a.s. and the residual fraction reaching deeper soil horizons. These parameters are also involved in several existing groundwater leaching indices such as the GUS (Groundwater Ubiquity Score; (Gustafson, 1989)), the LIX (Leaching Index; (Spadotto, 2002)), the RLPI (Relative Leaching Potential Index; (Hornsby et al., 1993)), and the AF (Attenuation Factor; (Rao et al., 1985)). As illustrated in Figure 5, the transfer potential to groundwater decreases sharply with increasing sorption capacity. Active substances characterized by low K_OC_ values exhibit the highest leaching potential, particularly when soil DT_50_ is high. Indeed, poorly persistent a.s. tend to degrade before reaching GW and strongly sorbed a.s. remain mostly retained in the upper soil layers, limiting their transfer.

Finally, aqueous solubility can influence the leaching behavior of a.s. Poorly soluble compounds tend to be retained in the soil, whereas those with high solubility are prone to leaching toward groundwater. However, several studies have demonstrated that this influence remains limited in environmental conditions, as concentrations of a.s. measured in soils after their application are generally below their solubility (Gustafson, 1989; Grill et al., 1999; Delvin et al., 2008). Hence, this parameter was not used in the Phytorisque model.

**Within the non-cultivated terrestrial compartment**, exposure of non-target organisms depends on two main pathways, as reported by several authors (Kumar et al., 2021; Authority (EFSA) et al., 2022).

On the one hand, exposure occurs through direct contact with the a.s. intercepted by the crop canopy and thus available to organisms moving across the crop. On the other hand, exposure results from the fraction dispersed beyond treated areas (through erosion, volatilization, and spray drift), thereby affecting the environmental habitats located near treated crops. The relative importance of these two exposure pathways on the five studied a.s., is illustrated in Figure 6. For most of them, exposure is largely dominated by direct contact within treated areas, whereas off-site dispersion represents a secondary pathway. However, clomazone present a different behaviour, as dispersion processes contribute substantially to its mobility. This highlights the importance of considering both mechanisms, as each may prevail depending on the characteristics of a.s. In addition to mobility, the exposure is strongly influenced by the a.s. degradation kinetics in the soil (soil DT_50_) and by their volatilization (P_v, sat_) on the crop canopy.

**In cultivated soils**, the duration of exposure of soil biota to an a.s. primarily depends on its soil DT_50_ as reported by several authors (Loose et al., 2025; Chandra et al., 2026). However, persistence within biologically active layers may be moderately limited by leaching, which reduces the residence time of the a.s. in the upper soil layer. Consequently, the soil adsorption coefficient (K_OC_) indirectly affects the effective exposure duration of soil biota by controlling the infiltration rate. This interaction between persistence and sorption is illustrated in Figure 7: while the soil DT₅₀ governs the initial increase in persistence, higher K_OC_ values increase persistence factors by limiting soil infiltration and retaining substances within the upper soil layers. Several authors have also shown that the persistence of a.s. in soil increases with sorption and degradation coefficients (Wauchope et al., 2002; Kah et al., 2007).

Finally, although exposure is important for assessing the potential effects of a.s. in the environment, it should be considered together with their intrinsic hazard to estimate overall risk, as implemented in the Phytorisque model. For example, at equivalent exposure, lambda-cyhalothrin is estimated to be significantly more hazardous than bromuconazole, which explains its particularly high Global Risk Index despite the relatively small quantities used in the Wallonia region (Table 3).

### 4.3. Effects of environmental parameters on transfer processes

Although the properties of a.s. influence ecosystem exposure, other factors including pedological, hydrological, and meteorological conditions in the area where these a.s. are used, also play an important role. The influence of annual climatic conditions on potential transfer of a.s. to surface waters is shown in Figure 2 for 2021 and 2022, the wet and dry years, respectively. A significant difference in the transfer potential to surface waters was observed between the two years. Humid conditions are associated with higher transfer potential compared to dry conditions. Additionally, the tested a.s. are transferred to water mainly in the dissolved phase, except for lambda-cyhalothrin, which is predominantly transported in the particulate phase. These findings highlight the importance of considering both the physicochemical properties of a.s. and the prevailing meteorological and pedological conditions when evaluating their transfer to water, and thus their potential contribution to the exposure of environmental compartments.

Overall, environmental conditions have two effects on the transfer models. On the one hand, they impact directly on the importance of the fluxes considered (e.g. importance of rainfall on runoff or erosion). On the other hand, environmental conditions will also affect the sensitivity of the model to a.s. physico-chemical properties. For instance, transfer to SW is less sensitive to soil DT_50_ during wet years, as the recurrence of rainfall events increases the probability of an a.s. to be transferred to this compartment before its degradation.

## 5. CONCLUSION

The assessment of environmental risk associated with the use of PPPs relies on the ability of indicators to accurately reflect the risk dynamics. The Phytorisque model is built on the evaluation of the exposure-toxicity ratio, in contrast to scoring indicators that simplify parameters without maintaining proportionality among risk components (Whiteside et al., 2008; Feola et al., 2011; Meys et al., 2024; Obregon et al., 2025).

Maintaining this proportionality is essential when objectives are expressed as percentage reductions in risk, ensuring that observed variations genuinely reflect real and comparable changes in risk levels.

Owing to its multi-compartment structure, the Phytorisque model provides an integrated view of risk by considering all environmental compartments potentially affected, which therefore ensures a coherent and complete assessment of risk at the ecosystem scale.

Currently, the model does not account for certain processes and dimensions of risk, such as degradation metabolites, preferential flow pathways in soils, or chronic toxicity effects, or even combined effects of pesticide applications (EFSA Scientific Committee et al., 2019). Furthermore, the parameterization of the current version of the model is not adapted to specific crops such as orchards.

Nevertheless, these limitations do not undermine the relevance of the tool within its current scope of application, and the model is designed as an evolving framework intended to be further refined and expanded. Phytorisque therefore represents a robust decision-support tool for temporal risk monitoring at the regional scale, for identifying the a.s. or sectors contributing most significantly to the global risk, and for the prospective evaluation of scenarios, whether comparing crop management practices or exploring strategies for the substitution of a.s. Moreover, the model is transferable to other regions through the adaptation of parameters to local pedoclimatic and land-use conditions, and the methodology can be adjusted where necessary if the local context differs significantly.

## Acknowledgement

Corder ASBL gratefully acknowledges the Public Service of Wallonia – Agriculture, Natural Resources and Environment, through the Directorate for Environmental Status and the Agriculture–Environment Integration Unit, for the full financial support of the project “Estimation quantitative des utilisations des produits phytopharmaceutiques par les différents secteurs d’activité en Wallonie” (grant number 03.02.02-25-78-73), and for the trust placed in our organization.

The research team also wishes to express its sincere gratitude to the members of the Steering Committee, whose involvement made it possible to successfully carry out the project missions through monitoring committees, working group meetings: Christine Cuvelier (SPW-DGARNE-DEMNA-DEE), Denis Godeaux (SPW-DGARNE-DEE-CIAE), Joëlle Vandersteen (Office of the Minister for the Environment), Olivier Miserque (SPW-DGARNE-DEMNA-DAEA), Pierre Nadin (FPS-HFCSE), Philippe Delaunois (SPW-DGARNE-DD-DRD) and Vincent Van Bol (FPS-HFCSE).

## Authors contributions

Conceptualization Loïc Monseur, Jean Baptiste de Maere, Chloé Guillitte, Laurence Janssens

**Data collection and analysis**: Loïc Monseur, Jean Baptiste de Maere, Chloé Guillitte

**Methodology**: Loïc Monseur, Jean Baptiste de Maere; Chloé Guillitte

**Writing original draft**: Loïc Monseur, Jean Baptiste de Maere; Chloé Guillitte

**Writing-review and editing**: Loïc Monseur, Jean Baptiste de Maere; Chloé Guillitte, Gaspard Nihorimbere, Laurence Janssens, Claude Bragard

**Funding acquisition:** Laurence Janssens & Claude Bragard

**Project Administration:** Laurence Janssens & Claude Bragard.

